# On improving experimental binding affinity predictions with synthetic data

**DOI:** 10.64898/2026.03.02.708607

**Authors:** Kevin Ryczko, Phyo Phyo Zin, Jordan Crivelli-Decker, Ly Le, Punit K. Jha, Harish Ramadas, Lorenzo Boninsegna, Justin Overhulse, Benjamin J. Shields, Pablo Lemos, Sasaank Bandi, Maarten van Damme, Amogh Sood, Lee Huntington, Mary Pitman, Martin Ganahl, Andrea Bortolato

**Author notes:** All work was done at SandboxAQ (Sasaank Bandi).

## Abstract

The success of deep learning binding affinity prediction models depends critically on expanding experimental data with reliable synthetic data. We extend the Structurally Augmented IC50 Repository (SAIR) with ≈80K absolute free energy perturbation (AFEP) calculations and present two distinct data splits, SAIR-FEP and SAIR-OOD (out-of-distribution), to simulate realistic drug discovery scenarios. We compare sequence-based proteochemometric (PCM) models and state-of-the-art, structure-based deep learning models and demonstrate that PCM models can be enhanced by physics-based descriptors in the cross-target hit-identification setting. While structure-based deep learning methods capture finer geometric detail, their performance is highly sensitive to the input structure. By filtering for high-confidence, co-folded complexes, we show that the performance improves predictably, whereas training on all complexes blindly limits performance gains. Finally, using the SAIR-OOD split, we demonstrate that simultaneous training on synthetic and experimental data improves performance on publicly available, experimental benchmarks. For the OpenBind benchmark, our models yield the lowest root mean squared errors and a 1.85x improvement of Spearman ranking correlation when compared to the original AEVPLIG model training on experimental data alone. These results provide a clear strategy for using synthetic data to advance experimental binding affinity predictions.

## 1 Introduction

Predicting the binding affinity between small molecules and their protein targets is a fundamental task in drug discovery, guiding both hit identification and lead optimization. While experimental techniques provide accurate affinity measurements, they remain resource-intensive and therefore cannot evaluate the vast chemical and proteomic spaces relevant to modern therapeutic design [1, 2]. To address these challenges, physics-based and machine-learning methods have become essential for prioritizing candidate compounds and accelerating drug discovery.

Historically, structure-based approaches such as molecular docking, scoring functions, and physics-informed simulations have served as the primary computational tools for estimating binding strength [3–5]. Although widely used, these methods often struggle with limited accuracy, complex parametrization, imperfect energy approximations, and challenges in handling protein flexibility [6]. More recently, deep learning (DL) models have gained momentum by learning nonlinear relationships directly from biochemical data. Models leveraging 3D protein–ligand structures, graph neural networks (GNNs), or molecular descriptors have demonstrated improved predictive performance [7–10], yet they remain heavily dependent on curated training data and carefully engineered molecular representations in order to be effective.

Structure-based deep learning models leverage 3D information from protein–ligand complexes to learn interaction patterns that correlate with binding strength. Early successes include voxelized 3D convolutional neural networks such as K_DEEP_ [7, 11, 12] and atomic convolutional networks [9], which demonstrated that CNNs can capture spatially local interactions in binding pockets. GNN architectures have also achieved strong performance by encoding complexes as interaction graphs; recent examples include AEVPLIG [13] and GEMS [14], both of which operate on a ligand graph, and [15], which fuses protein and ligand graphs together. Hybrid physics–ML methods further integrate physics-based features with learned representations to improve generalization [16].

Despite these advances, structure-based models rely heavily on large quantities of high-quality structural data and accurate binding affinity data. While open-access resources exist (such as the RCSB Protein Data Bank [17], and PDBBind [18]), these sources are often imperfect and contain significant biases that may undermine models trained on them. For example, several analyses have demonstrated that deep networks may exploit biases in datasets such as PDBbind [18], often memorizing instead of learning true intermolecular interactions [19]. Recent advances in computational structure-prediction approaches have now made it possible to generate large amounts of accurate structural data to augment existing datasets or even to predict the bound conformation of a protein-ligand complex [20]. This, in theory, should enable the generation of additional high-quality structural data for training, thereby improving the performance of structure-based models. However, recent work [21] shows that naively training on this co-folded data does not yield more performant models, with predictive performance failing to improve as the amount of added data increases. Thus, it remains unclear which approach is most effective for integrating synthetic data into the training sets of DL affinity models.

To overcome limitations in structural data, one can turn to Simplified Molecular Input Line Entry System (SMILES) and sequence-driven approaches, such as proteochemometric (PCM) models. Models such as ChemBoost [22] and WideDTA [23] treat protein sequences and SMILES strings as textual inputs, learning “chemical language” embeddings to predict affinity. More recent models integrate graph-based representations with sequence features, such as GNNSeq [24], which combines graph neural networks with sequence-derived descriptors to achieve competitive performance without requiring protein–ligand complex structures. However, binding affinity depends critically on structural information, so structure-based models are expected to better capture the underlying physics, leading to better generalization.

Overall, the literature illustrates a diverse landscape of affinity prediction methods, each with different assumptions about data availability and molecular representation. Structure-based DL offers fine-grained geometric information, while language-based approaches provide flexibility and broad applicability. However, systematic comparisons under controlled conditions remain limited in the literature thus far. In addition, regarding structure-based approaches, it remains unclear what the best practices are for training a model on synthetic structures. This motivates our benchmarking study, which evaluates language-model–based predictors against DL and classical structure-based approaches using consistent datasets, metrics, and evaluation protocols.

In this work, we train PCM and structure-based models (AEVPLIG) on the Structurally Augmented IC50 Repository (SAIR) dataset [25], evaluating the impact of including augmented structural data during training and comparing their performance in predicting experimentally derived binding affinity data. We first compare them directly on a subset of SAIR, called SAIR-FEP, where we’ve computed 80K absolute free energy perturbation (AFEP) calculations along with various docking scores. We investigate the performance of PCM models using multiple permutations of physics-informed features to identify the most effective feature-stacking strategies for binding affinity prediction. Next, we study the performance of structure-based models on publicly available benchmarks when trained on both experimental and synthetic data simultaneously. To do so, we create another subset of SAIR called SAIR-OOD (out-of-distribution), in which all complexes that overlap with the experimental complexes were removed to avoid data leakage (Figure 1). We aim to understand model performance across various splits and to probe the underlying data to identify best practices for machine learning applied to synthetic data for binding affinity prediction.

**Fig. 1.**
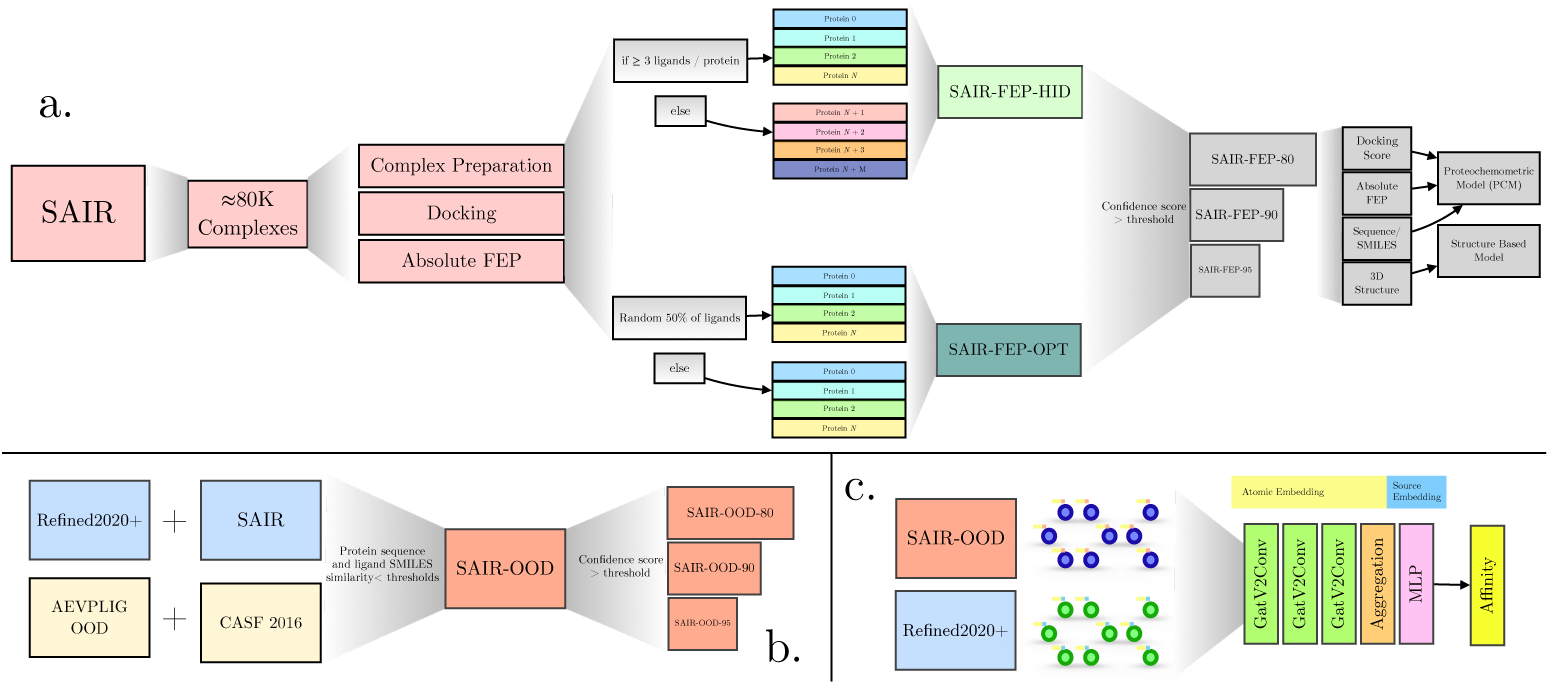
Overview of the work presented. a) The construction of the SAIR-FEP dataset, which includes computed docking and absolute free energies for *≈*80K complexes. Hit identification (SAIR-FEP-ID) and lead optimization (SAIR-FEP-OPT) splits are done to evaluate models on different drug discovery tasks. The data is then further filtered using the co-folding confidence score and then used for training downstream models. b) The construction of the SAIR-OOD dataset, which requires filtering out complexes that have a high protein or ligand similarity with test sets evaluated (AEVPLIG-OOD and CASF 2016). This dataset is then further filtered using the co-folding confidence score. c) Depiction of the AEVPLIG model with the additional source embedding, allowing for training on synthetic and experimental complexes simultaneously.

## 2 Methods

### 2.1 Curation process of SAIR-FEP

#### 2.1.1 The SAIR Dataset

The SAIR dataset [25] consists of predicted co-folded structures generated from Boltz-1x [20] of 5,244,285 structures across 1,048,857 protein-ligand complexes sourced from ChEMBL [26, 27] and BindingDB [28, 29]. As described in [25], the structures were produced with the Boltz-1x release using three recycling steps and 200 sampling steps (default settings otherwise), with multiple sequence alignments from MMseqs2 (UniRef30 v2302 and ColabFoldDB v202108). This structural data is also paired with experimentally derived binding data to train binding affinity models.

#### 2.1.2 Filtering of SAIR

In this work, we filter the SAIR dataset to ensure the highest quality structures are used in our analyses. For each complex, 5 conformations were generated with Boltz-1x, with multiple entries possible for certain complexes due to assay variability in IC50 measurements. When a protein–ligand pair had more than one reported affinity measurement, we did not unify these into a single label; each measurement was retained as a separate training entry, so the model is trained on the repeated complex paired with each of its experimental values, preserving the natural assay-to-assay variability rather than imposing an arbitrary aggregation. We selected only the top-ranked structure for each protein-ligand complex with an aggregated confidence score (estimating reliability via the predicted local distance difference test, pLDDT, and interface predicted TM-score, ipTM) greater than 0.5, yielding 1,754,697 complexes. These were further filtered to retain only biochemical assays, proteins with at least 10 measurements, and those with sufficient pIC50 variability ( 2.5 log units). The filtered data had 1,641 unique proteins and 100,617 entries. Next, we focus on curating 2 train/test splits that emulate realistic drug discovery scenarios. SAIR-FEP-HID (Hit Identification): This split evaluates zero-shot model generalization across unseen protein targets. We partitioned the data protein-wise: high-count proteins ( 3 ligands) were randomly allocated to 70% for training and 30% for testing. To maximize the challenge of target generalization, all remaining low-count proteins (*<*3 ligands) were exclusively assigned to the test set, ensuring the model is evaluated on rare targets. SAIR-FEP-OPT (Lead Optimization): This split evaluates model performance when proteins are shared but ligands are split randomly between the training and test sets. This emulates a lead optimization campaign, where the target and some hit molecules are known, and the model must identify better molecules against established hits. Dataset statistics for these splits are shown in Table 1.

**Table 1.** Details of training and test sets used in this report. The values 80/85/90/95 indicate the confidence value (of the co-folding model) threshold applied. For example, SAIR-OOD-90 indicates that a confidence score threshold of 0.9 was applied. *N*_proteins_ counts distinct protein sequences; for CASF 2016 this is larger than the number of core-set target clusters (49 of the 57 PDBbind/CASF clusters are represented) because a single target includes several point mutants or constructs with near-identical sequences (see the per-target analysis in the Supplementary Information).

| Name | *N*<sub>proteins</sub> | *N*<sub>ligands</sub> | *N*<sub>complexes</sub> |
|---|---|---|---|
| SAIR | 5149 | 734359 | 1043300 |
| SAIR-FEP-HID<sub>train</sub> | 1107 | 48461 | 55083 |
| SAIR-FEP-HID<sub>test</sub> | 505 | 22053 | 23519 |
| SAIR-FEP-HID-80<sub>train</sub> | 818 | 29765 | 32918 |
| SAIR-FEP-HID-85<sub>train</sub> | 609 | 20019 | 21775 |
| SAIR-FEP-HID-90<sub>train</sub> | 319 | 8571 | 9195 |
| SAIR-FEP-HID-90<sub>test</sub> | 152 | 3895 | 4015 |
| SAIR-FEP-HID-95<sub>train</sub> | 89 | 1435 | 1480 |
| SAIR-FEP-HID-95<sub>test</sub> | 43 | 849 | 863 |
| SAIR-FEP-OPT<sub>train</sub> | 1223 | 33762 | 36880 |
| SAIR-FEP-OPT<sub>test</sub> | 1223 | 34358 | 37506 |
| SAIR-OOD | 5089 | 723555 | 1031912 |
| SAIR-OOD-80 | 3219 | 399054 | 536118 |
| SAIR-OOD-85 | 2297 | 261812 | 339582 |
| SAIR-OOD-90 | 1222 | 102144 | 120614 |
| SAIR-OOD-95 | 285 | 15940 | 17076 |
| Refined2020+ | 11130 | 14400 | 18022 |
| Refined2020++ | 4822 | 7294 | 8240 |
| AEVPLIG-OOD | 189 | 277 | 291 |
| CASF 2016 | 156 | 193 | 193 |

#### 2.1.3 Complex Preparation

Co-folded protein-ligand structures were initially prepared by separating the ligand and protein into their apo forms. Next, ligands were processed with RDKit [30] to ensure correct protonation states, tautomer assignment, and to resolve stereochemistry. The binding poses generated by co-folding were reserved for Gnina scoring [12] and AFEP calculations [31]. Proteins were prepared to ensure proper protonation of key residues, repair missing atoms or side chains, and perform short energy minimizations to resolve steric clashes or unrealistic geometries.

#### 2.1.4 Docking Rescoring Using CNN and Vinardo Scoring Functions

Prior to absolute binding free energy calculations, all complexes were rescored using Gnina [12] with both a convolutional neural network (CNN) based scoring function and the Vinardo empirical scoring function [32]. This rescoring step served as an intermediate triage layer to benchmark machine-learning-based and classical docking scores on the same physics-enriched structural ensemble and to enable direct comparison with subsequent AFEP-derived binding free energies. The CNN evaluates protein–ligand complexes represented as three-dimensional voxel grids to predict binding affinity and pose quality directly from learned spatial interaction patterns, while Vinardo is a physics-inspired empirical scoring function derived from AutoDock Vina [33] that estimates binding affinity as a weighted sum of intermolecular interaction terms optimized for robustness and transferability.

#### 2.1.5 Free energy calculations

For each protein-ligand complex, free energy calculations were performed using the AQFEP protocol described in [31]. AQFEP uses an absolute free energy perturbation calculation based on the double-decoupling alchemical protocol. The double-decoupling approach is considered the “gold standard” for absolute free-energy calculations by ensuring thermodynamic consistency, accurate sampling of the free-energy landscape, and broad applicability across a variety of systems. Using an absolute free energy calculation, which directly estimates the binding free energy of the given ligand-protein pair, the procedure requires much less human guidance than the more typical relative free energy calculations. Unlike relative FEP, which requires a congeneric ligand series and predefined alchemical mappings between similar compounds, AQFEP is an absolute FEP method that evaluates each ligand independently, making it directly applicable to heterogeneous libraries spanning diverse chemotypes and binding modes. This independence enables more flexible benchmarking across targets and chemistries, and facilitates systematic comparison with docking and machine–learning–based scoring methods in large-scale virtual screening workflows. Convergence was assessed independently for both the bound and unbound legs using predefined criteria, and only complexes satisfying all convergence requirements were retained for downstream analysis. Binding free energies are reported as Δ*G* (kcal/mol), with associated uncertainties reflecting the statistical error of the free energy estimates. In total, roughly 157,000 GPU hours (NVIDIA Tesla T4) were used for the FEP calculations.

### 2.2 Curation of SAIR-OOD

As mentioned in previous literature [19], benchmarks on structure-based affinity models have revealed biases in the underlying training data, highlighting data leakage and the need for proper held-out test sets to rigorously evaluate these models. To address this, we evaluated the impact of training on two datasets: Refined2020+ [13] (which is a specialized split of PDBBind2020 [18]) and SAIR [25]. To better understand our model’s generalization performance and the impact of synthetic data, we evaluated on two commonly used test sets in the literature: AEVPLIG-OOD [13], and CASF 2016 [34]. However, initial analyses revealed high structural similarity between our training sets and the two test sets. As a result, we curated a second dataset, SAIR-OOD, that addresses this limitation. The curation approach is described below.

To minimize data leakage, we removed entries from the training set that had high structural or sequence similarity to the evaluation sets, as done in previous work [13]. To calculate protein similarity, we used MMSeqs2 [35], and to calculate ligand similarity, we used Tanimoto similarity between entries’ Morgan fingerprints, as included in RDKit version 2025.09.03 [30]. If the protein or the ligand had a similarity value *>* 0.5, the complex was removed from the training set. With SAIR, 5 complexes were predicted per complex. However, since structure-based models require a single structure, we chose to use the complex with the lowest Vina score [4]; this left 1,043,300 unique structures. Filtering these complexes based on sequence and SMILES similarity left 1,031,912 structures. In addition, we further filtered this set based on the confidence scores output by Boltz-1, which are included in the SAIR dataset. We considered confidence values of 0.8, 0.85, 0.9, and 0.95, yielding 4 unique training datasets: SAIR-OOD-80/85/90/95. The number of entries, along with the unique number of proteins and ligands, is shown in Table 1.

In addition to the filtering of SAIR, we also filtered the entries of the Refined2020+ [13] to eliminate overlap with the CASF 2016 [34] test set. We refer to this training data as Refined2020++.

### 2.3 Training of Proteochemometric Models

We developed a Proteochemometric (PCM) modelling framework to predict the binding affinity (potency) of small molecules against multiple protein targets. This approach utilizes a supervised regression architecture that simultaneously learns from ligand chemical space and protein sequence space.

#### 2.3.1 Feature Representation

Molecules were represented as SMILES strings and featurized using Morgan fingerprints (2048 bits, count-based). Protein targets were represented by their primary amino acid sequences. To capture high-dimensional biological context, we employed pre-computed embeddings from the ESM-2 (Evolutionary Scale Modeling) 650M-parameter model (esm2 t33 650M UR50D) [36]. The 1280-dimensional per-residue embeddings were reduced to 320 dimensions with a fixed random (Johnson–Lindenstrauss) projection and mean-pooled over residues to yield a single 320-dimensional descriptor per protein, providing a dense representation of protein sequences that captures evolutionary and structural information. Physics-based features: To evaluate the contribution of physics-based descriptors, we systematically trained models with docking scores, including minimized affinities (the empirical Vinardo binding affinity reported by gnina after local energy minimization of the cofolded pose) and deep-learning-based scoring (CNNscore, CNNaffinity) from gnina [12] as well as free energy of binding and solvation descriptors (from AQFEP).

#### 2.3.2 Model Architecture and Optimization

PCM models were implemented with Pytorch [37] using an ensemble approach that included Gaussian Mixture Network (GMN) [38], Mean Variance Estimation (MVE), and Random Forest (RF). Hyperparameter selection was conducted via an automated Bayesian optimization framework using cross-validation. The search space encompassed critical parameters, including feature bit counts and network layer dimensions, while the preprocessor block remained frozen to maintain feature consistency. The optimization process was initialized with an *n* = 1 random sampling phase, followed by 9 iterations of Bayesian search directed by a surrogate probability model. The objective function was set to maximize the Spearman correlation coefficient. Upon completion, the configuration yielding the best validation performance was selected, and the final models were refitted on the full training dataset to maximize data utility and stability.

### 2.4 Training of structure-based models

The structure-based model investigated in this work is a GNN called AEVPLIG [13] which operates on molecular graphs. Briefly, the nodes and edges in the graph represent the heavy atoms of the ligand. The heavy atoms of the protein pocket are used to construct node-level embeddings but are not explicitly included in the graph passed to the GNN. An element of the embedding for heavy atom *i* from a particular ligand is given by

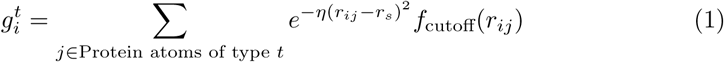

where

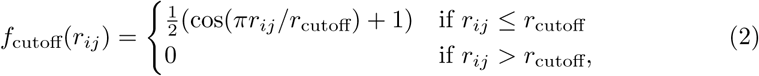

*r_ij_* is the radial distance between ligand atom *i* and protein pocket atom *j*. One defines a set of values for *η* and *r_s_* yielding a vector embedding for the set of protein atoms of type *t*. All of the different protein atoms are then concatenated to form the full input vector, **g_i_**. When training on synthetic and experimental data simultaneously, there is an additional embedding vector **s** that is concatenated with the atomic input vectors **g_i_**. This embedding represents the source of the data and the structure it is derived from, and can take 2 forms: synthetic or experimental.

We train the models for 200 epochs using a batch size of 128, a hidden dimension of 256, 3 attention heads, a learning rate of 0.0001229, and the leaky ReLU activation functions throughout. The ensemble model consisted of 5 models, all of which were trained using different seeds of the same data. We refer the curious reader to the original manuscript for more details [13]. When training on synthetic and experimental data simultaneously, an additional embedding vector is concatenated with the atomic input vectors. This embedding represents the source of the data and the structure it is derived from, and can take 2 forms: synthetic or experimental.

## 3 Results

### 3.1 Model performance on SAIR-FEP

We first characterise model behaviour on the SAIR-FEP data, whose experimental affinity labels are paired with co-folded structures, to establish the conditions under which augmented structural data is beneficial, namely, which physics-based descriptors improve feature-based models and how the quality of the co-folded structures governs performance. These conditions are then exploited in Section 3.2, where joint training on synthetic and experimental data is evaluated directly on experimental benchmarks.

#### 3.1.1 Including docking and FEP results improves predictions from proteochemometric models

To establish baseline performance for feature-based models, we first evaluated the PCM model using the SAIR-FEP-HID train/test split. As mentioned in Section 2, an array of physics-based scoring methods was included in this split to evaluate if including these metrics improves model performance. These values were appended to the PCM model’s existing features. We compare this model to a state-of-the-art structure-based model, AEVPLIG, without additional physics-based features included during training. As shown in Figure 2a, we observe improvements across all metrics when including physics-based scores as features. 3 of 4 models that included docking scores (outputs of gnina docking, including those from the convolutional neural network (CNN)) all improved over the baseline model. However, because some gnina metrics depend on the CNN model’s output, data leakage may occur, potentially leading to results that do not generalize. To investigate which gains are genuinely physics-derived, Figure 2a separates the CNN-derived docking scores from the purely physics-based (Vinardo and AFEP) descriptors: the physics-only PCM + AFEP model improves only modestly over the PCM baseline (Pearson 0.420 versus 0.405), whereas the largest gains arise from the CNN-derived docking scores (PCM + Docking, Pearson 0.461). The latter is likely partially inflated by circularity: the gnina CNN was itself trained on protein–ligand complexes that overlap with our evaluation data. Quantifying this against PDBbind (the basis of gnina’s CrossDocked2020 CNN training set), 25.3% of the SAIR-FEP-HID test complexes contain a protein with 95% sequence identity to a PDBbind protein and 20.2% contain a ligand with ECFP4 Tanimoto 0.5 to a PDBbind ligand, so the CNN score can carry information about the test labels. By contrast, the AFEP and Vinardo descriptors are computed de novo for each complex and do not share training data with the test set. In some cases, combining different scoring functions (e.g., PCM + Docking + AFEP) yields worse metrics than using them independently (e.g., PCM + Docking or PCM + AFEP). This indicates that including multiple scoring functions could introduce contradictions, inevitably leading to poor model performance. AEVPLIG outperformed most PCM models and was on par with the best-performing PCM model, including docking scores, indicating that the non-linear mappings learned directly from structural data are sufficient to produce a high-quality model. Interestingly, including physics-based docking and AFEP features in the node embeddings of AEVPLIG models did not improve performance on the test set.

**Fig. 2.**
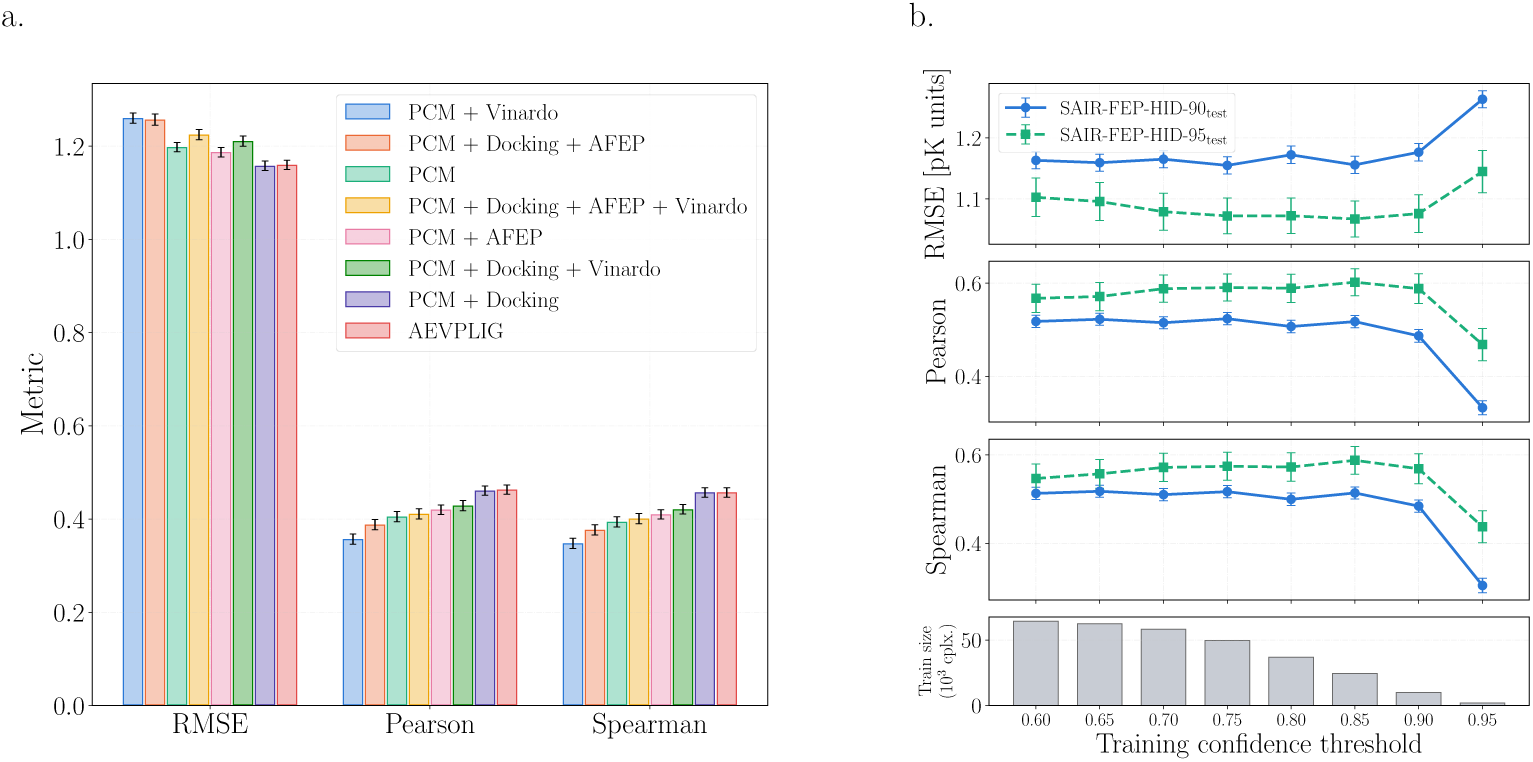
a) Metrics of PCM with the addition of docking and absolute free energy perturbation values as input features, as well as AEVPLIG evaluated on the SAIR-FEP-HID test set. Physics-based data enhances PCM predictions. b) Top three panels: metrics (RMSE, Pearson, Spearman) tracked across the SAIR-FEP-HID-90/95test sets for AEVPLIG models trained at co-folding confidence thresholds spanning 0.60–0.95; error bars are *±*1 s.d. from a 1000-iteration bootstrap of the ensemble predictions. Bottom panel: the corresponding number of training complexes, which grows roughly 35*×* (from *≈* 1.9 *×* 10^3^ at threshold 0.95 to *≈* 6.4 *×* 10^4^ at 0.60) as the filter is relaxed. Despite this large increase in data, performance is essentially flat across the broad 0.60–0.85 range and degrades sharply only at the most aggressive filters (0.90–0.95), where too few high-confidence complexes remain to train on, indicating a near-optimal plateau of the dataset-size versus structural-quality trade-off.

#### 3.1.2 Quality of co-folded structures impacts downstream model performance

Our initial analysis revealed that AEVPLIG was among the best-performing models on the SAIR-FEP-HID split. We note, however, that this comparison varies both the model architecture and its inputs, a structure-based network trained on co-folded complexes versus feature-based PCM models, and therefore does not, on its own, isolate the contribution of the augmented structural data from that of the structure-based model. The specific effect of augmentation is instead quantified in Figure 4a, where the same AEVPLIG architecture is trained with and without synthetic data on identical experimental data. Motivated by the strong performance of the structure-based model, we now consider the effects of the quality of the underlying structural data used during model training. To start, we filter SAIR-FEP-HID with confidence scores ranging from 0.60 to 0.95 in increments of 0.05, each yielding its own train/test split. Confidence scores were obtained from the pLDDT head of the Boltz-1x model. This was introduced for alphafold models and has been shown to correlate well with binding poses [39, 40]. We then monitor the performance of AEVPLIG models on the 0.9 and 0.95 confidence-score test splits (called SAIR-FEP-HID-90_test_ and SAIR-FEP-HID-95_test_, respectively) as we add more data with lower confidence scores. As shown in Figure 2b, we observe saturation of all metrics as we include more data, which is contrary to scaling law behaviour that is typically observed in deep learning models. An interesting observation is that across the broad 0.60–0.85 range the metrics are essentially flat, for SAIR-FEP-HID-95_test_ the Spearman correlation varies only between 0.55 and 0.59, peaking at the SAIR-FEP-HID-85_train_ split, and they degrade sharply only at the most aggressive filters (0.90 and especially 0.95), where too few high-confidence complexes remain to train on. Notably, this plateau holds even though the training set grows by roughly 35 as the threshold is lowered (from 1,850 complexes at 0.95 to 64,400 at 0.60; bottom panel of Figure 2b). This suggests that a careful balance between dataset size and the quality of co-folded structures used during training affects model performance: over a wide range the competing effects of adding more data and retaining only higher-confidence structures approximately offset, so the operating point used here lies on a near-optimal plateau rather than at a sensitive edge.

**Fig. 3.**
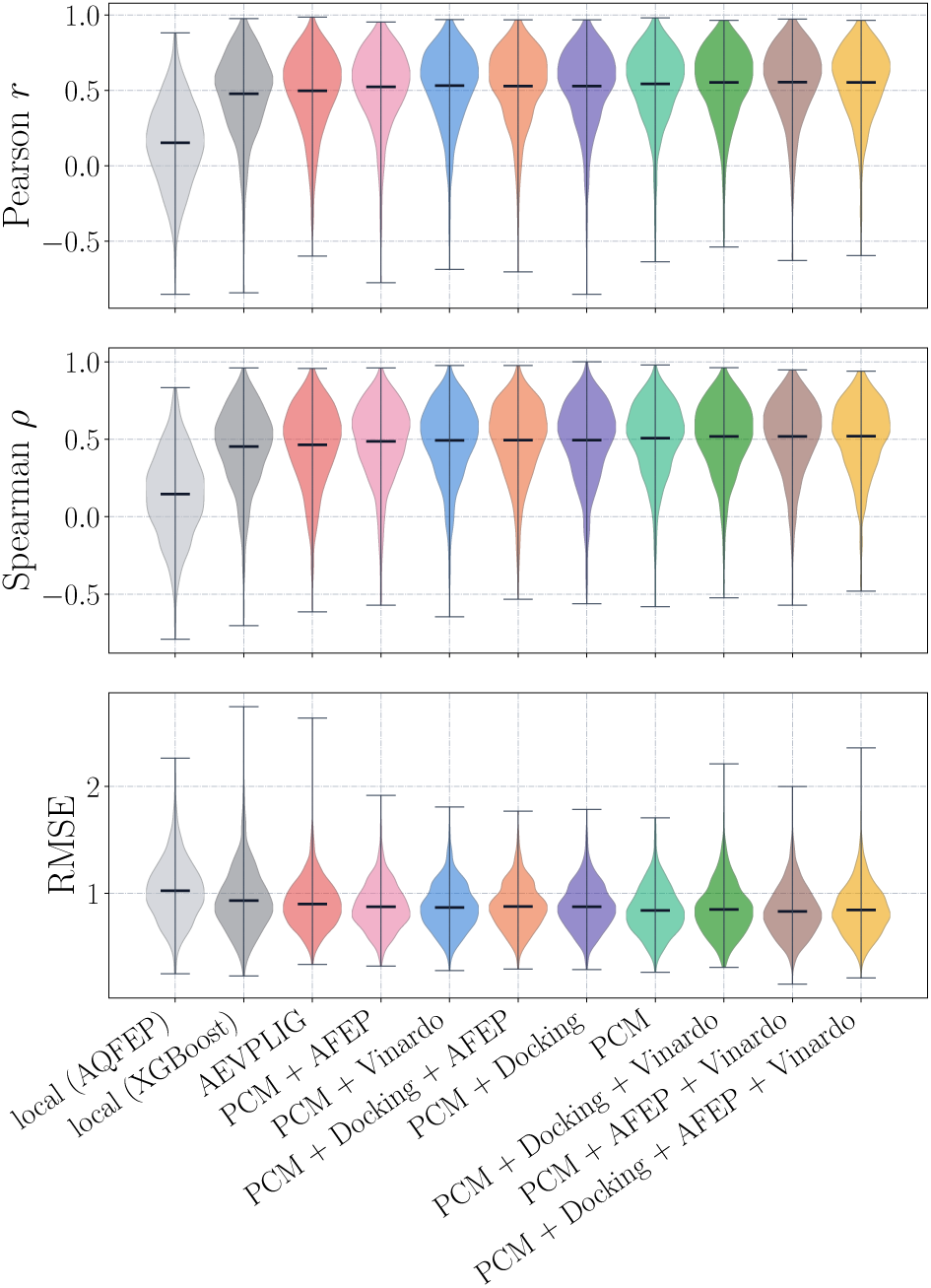
Comparison of Pearson/Spearman correlation coefficient and root mean squared error (RMSE) distributions across model architectures and feature integration strategies for the SAIR-FEP-OPT split for proteins with *≥* 10 ligands. Distributions represent the performance across individual protein targets; as each violin shows the full distribution (median and range) across targets, it conveys the complete spread of per-target performance rather than a summary error bar. AFEP-Linear and XGBoost represent “local” models trained independently for each protein (shown in gray), whereas AEVPLIG and PCM-based models utilize “global” training across the entire dataset; each model keeps the same color as in Figure 2a. While AEVPLIG achieves the tightest distribution overall, the PCM baseline outperforms the variants augmented with additional physics-based scoring features.

**Fig. 4.**
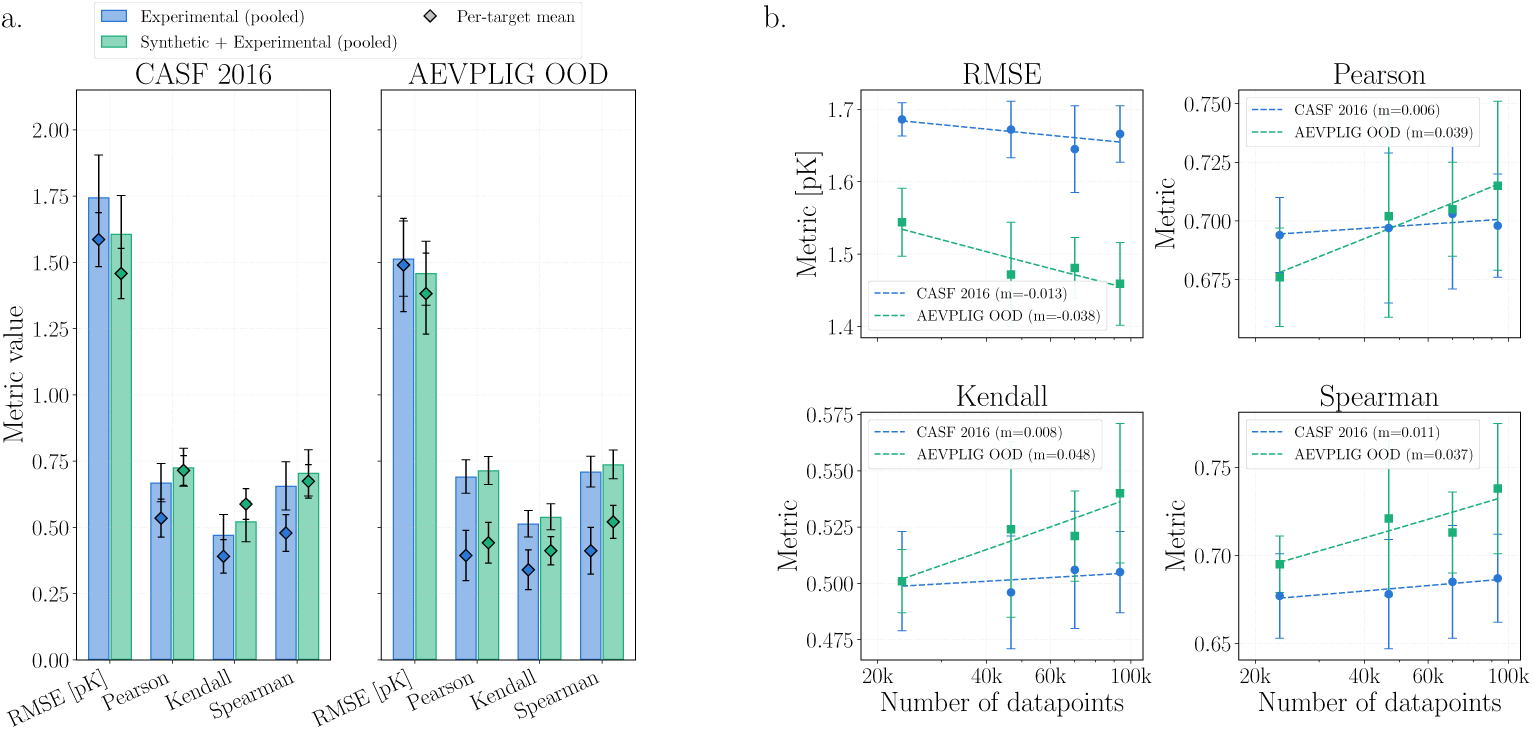
a) Test metrics comparing AEVPLIG models trained on experimental data alone versus models trained on synthetic and experimental data simultaneously. Consistent performance increases are seen when training on synthetic data. Bars show the pooled test metrics; overlaid diamonds show the per-target mean *±* standard error across targets with *≥* 3 complexes (Table 2). b) Scaling curves for various metrics versus the number of high-quality complexes (confidence score *>* 0.9) being trained on. Traditional scaling laws hold, with model performance consistently improving as dataset size increases.

**Table 2.** Per-target performance of the experimental-only baseline versus the SAIR-augmented model, averaged across targets (*±* standard error), with paired Wilcoxon signed-rank *p*-values. CASF 2016 targets are the PDBbind/CASF 2016 core-set clusters (90% sequence similarity); AEVPLIG-OOD targets are distinct protein sequences. Only targets with at least three complexes are included.

| Test set | Metric | Baseline | +SAIR | *p*-value |
|---|---|---|---|---|
| CASF 2016 (*n*=38) | RMSE [pK] ↓ | 1.586 ± 0.102 | 1.458 ± 0.095 | 0.006 |
|  | Pearson ↑ | 0.535 ± 0.072 | 0.715 ± 0.056 | 0.002 |
|  | Kendall ↑ | 0.391 ± 0.063 | 0.588 ± 0.058 | 0.008 |
|  | Spearman ↑ | 0.479 ± 0.069 | 0.674 ± 0.063 | 0.007 |
| AEVPLIG-OOD (*n*=22) | RMSE [pK] ↓ | 1.490 ± 0.176 | 1.382 ± 0.153 | 0.069 |
|  | Pearson ↑ | 0.394 ± 0.095 | 0.442 ± 0.077 | 0.160 |
|  | Kendall ↑ | 0.340 ± 0.075 | 0.412 ± 0.053 | 0.112 |
|  | Spearman ↑ | 0.412 ± 0.088 | 0.521 ± 0.062 | 0.086 |

#### 3.1.3 Model performance in a lead optimization scenario

We now consider the SAIR-FEP-OPT data split, in which proteins are shared across the training and testing sets, but ligands are split 50/50. As discussed in Section 2, this aims to test model performance in cases where the target and hit molecules are known, and to find new molecules that surpass current hits in binding affinity. Here, we consider independent models trained for each protein and a model trained across all proteins. For the per-protein models, we use linear regression with Δ*G* from the AFEP calculations as the only input, serving as the physics-based baseline. In addition, we train XGBoost [41] models on Morgan fingerprints with 1024 bits and radius of 2, computed with RDKit [30]. For all XGBoost models, we use a max tree depth of 4, 100 models, and a learning rate of 0.1. For the all-protein model, we use PCM variants as described in Section 3.1.1 and AEVPLIG. In all cases, we compute the Pearson and Spearman correlation coefficients and the root mean squared error (RMSE) for each protein with 10 or more ligands, and aggregate the coefficients across proteins. As shown in Figure 3, the physics-based linear regression model, relying solely on AFEP-derived Δ*G* values without data-driven calibration, naturally yields the lowest average correlation of 0.15 with a broad spread across targets. This behaviour is consistent with its sensitivity to small co-folding errors and the omission of protein reorganization and ligand entropic contributions in the fast AQFEP protocol, underscoring its value as a conservative baseline and the importance of thorough retrospective validation before prospective use. The PCM model, without additional physics-based features, was the top-performing architecture among those evaluated. Interestingly, the inclusion of physics-based metrics (Docking and AFEP) into the PCM variants resulted in a performance degradation compared to the base model. Although the mean correlations are 0.5 for the models, some individual targets show negative correlations, reflecting the intrinsic difficulty of within-target ranking for certain proteins. The AEVPLIG model barely surpasses the local XGBoost model in this case, suggesting that more complex structure-based models are not always the best choice.

The contrasting behaviour between the SAIR-FEP-HID and SAIR-FEP-OPT splits reflects how useful the physics-based descriptors are in each regime. Hit identification is a zero-shot, cross-target task in which the model must generalize to unseen proteins; there the absolute physics descriptors (docking and AFEP) carry complementary signal and improve the PCM models, and the pocket geometry captured by AEVPLIG is advantageous. Lead optimization is a within-target task in which the model has already seen the target during training. The physics descriptors rank only weakly within a target: across proteins with at least ten ligands, the median per-target ranking is *ρ* = 0.20 for AFEP Δ*G* and *ρ* = 0.26 for the gnina docking scores, well below the base PCM model’s within-target performance (*ρ* 0.5; Figure 3). This is not because the within-target affinity range is small (the median within-target range is 4 log units); rather, the fast AFEP and docking scores simply rank congeneric ligands poorly. Because the data-driven model already ranks the target’s ligands well, appending these weaker, noisier per-complex features dilutes its signal and degrades the PCM variants, and for the same reason the more expressive structure-based model offers little advantage over simple local models. Physics-based descriptors therefore enhance feature-based models specifically in the cross-target hit-identification regime, where the model cannot otherwise generalize.

### 3.2 Model performance on SAIR-OOD

#### 3.2.1 Training on synthetic data improves the performance of models on experimental benchmarks

Our previous analyses demonstrated that utilizing high-quality, structurally augmented data improves model performance on affinity prediction tasks. However, an open question in the literature is whether combining augmented data and experimentally derived structural data can improve model generalization performance. To address this, we allow the model to train on both experimental and synthetic data simultaneously and evaluate the model on two commonly used model generalization benchmarks used in the field. To accomplish this, we modified the AEVPLIG model architecture by introducing an additional embedding layer and adding an additional index to the data such that the correct embedding is selected during training and inference. This embedding was then concatenated to the node embeddings before applying the graph attention layers. A depiction of this is shown in Figure 1c. The PCM model requires no modification to train across synthetic and experimental data simultaneously. Although it performed well during training on Refined2020++ and testing on CASF 2016, we found that adding more data did not improve model performance. An initial baseline AEVPLIG model was trained on experimental data alone and compared to a model trained on both experimental and co-folded structures. All models were evaluated on CASF 2016 and AEVPLIG-OOD (see Section 2). As shown in Figure 4a, models trained on experimental data alone yield RMSEs from 1.5-1.7 pK units, and correlation statistics ranging from 0.51-0.71. Importantly, after adding synthetic data (SAIR-OOD-85/90) to the training set alongside experimental (Refined2020+/Refined2020++) data, we observe performance increases on both test sets. For CASF 2016, we observe an 8% decrease in the RMSE, and a 9-11% increase in the correlation metrics compared to the baseline. For AEVPLIG-OOD, we find a 4% decrease in the RMSE, and a 3-5% increase in the correlation metrics. Notably, although the SAIR augmentation data is composed entirely of IC_50_ measurements, CASF 2016 contains only *K_i_* (114 complexes) and *K_d_* (79 complexes) values; breaking the CASF 2016 comparison down by measurement type (Table 8) shows that augmentation improves both the *K_i_* (Pearson 0.668 0.733) and *K_d_* (Pearson 0.649 0.694) subsets, indicating that the synthetic data provides transferable structure–affinity signal rather than an IC_50_-specific label bias. Although the correlation increases may seem modest, they have significant performance increases in downstream drug discovery tasks. For example, consider the increase in hits identified during a virtual screening campaign. With a total library size of 1,000,000 and 1000 true hits, if the improvement from the Pearson correlation of 0.669 to 0.727, as shown in Table 3 for CASF 2016, the number of hits goes from 27 to 38 when filtering out the top 0.01% (1000 molecules). This is a 41% increase in the number of hits, which is a substantial gain even with a modest improvement in correlation. To confirm that these improvements are statistically significant rather than aggregate point estimates, we recomputed the metrics per target and compared the baseline and SAIR-augmented models across targets using a paired Wilcoxon signed-rank test (Table 2). On CASF 2016 (38 targets with 3 complexes), adding SAIR improves the per-target Pearson from 0.535 to 0.715 (*p* = 0.002), the Spearman from 0.479 to 0.674 (*p* = 0.007), and the RMSE from 1.586 to 1.458 (*p* = 0.006), all three significant. On AEVPLIG-OOD the per-target estimates are sparse (only 22 targets reach 3 complexes) and the two models are statistically indistinguishable (all *p >* 0.05), consistent with the already-strong baseline on that test set. This result suggests that including high-quality synthetic structural data improves the performance of structure-based affinity models on challenging out-of-distribution test sets. To verify that AEVPLIG exploits the 3D structure rather than memorizing ligand identity [19], we additionally ran a ligand-only baseline and a pocket-scrambled control (Table 9). On the ligand-disjoint AEVPLIG-OOD set, a ligand-only fingerprint model and a sequence+ligand PCM model both fall to Pearson 0.5, and scrambling the pocket (pairing each ligand with a random incorrect protein) degrades AEVPLIG from Pearson 0.708 to 0.402; only on the ligand-biased CASF 2016 set does the ligand-only model appear competitive, confirming that the structural information is genuinely used where generalization is required.

**Table 3.**
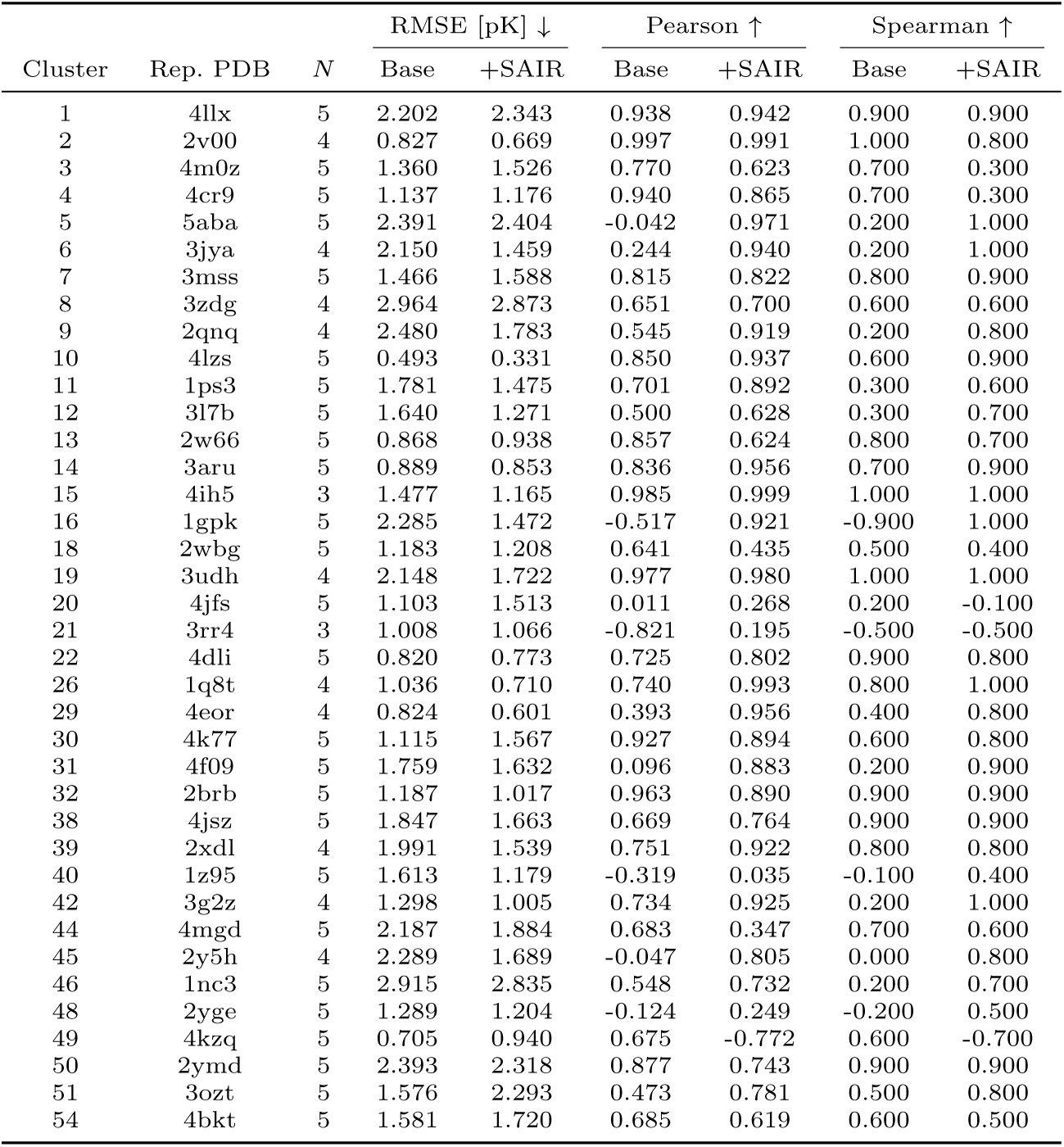
Per-target (CASF 2016 core-set cluster) assignments and paired performance underlying Table 2. Clusters are the PDBbind/CASF 2016 90%-sequence-similarity target clusters; the cluster index follows the core-set numbering (1–57). A representative PDB code is given for each cluster. Only the 38 clusters with at least three evaluated complexes are listed (176 complexes total). “Base” is the experimental-only model and “+SAIR” the synthetic-augmented model.

|  |  |  | RMSE [pK] ↓ |  | Pearson ↑ |  | Spearman ↑ |  |
|---|---|---|---|---|---|---|---|---|
| Cluster | Rep. PDB | *N* | Base | +SAIR | Base | +SAIR | Base | +SAIR |
| 1 | 4llx | 5 | 2.202 | 2.343 | 0.938 | 0.942 | 0.900 | 0.900 |
| 2 | 2v00 | 4 | 0.827 | 0.669 | 0.997 | 0.991 | 1.000 | 0.800 |
| 3 | 4m0z | 5 | 1.360 | 1.526 | 0.770 | 0.623 | 0.700 | 0.300 |
| 4 | 4cr9 | 5 | 1.137 | 1.176 | 0.940 | 0.865 | 0.700 | 0.300 |
| 5 | 5aba | 5 | 2.391 | 2.404 | -0.042 | 0.971 | 0.200 | 1.000 |
| 6 | 3jya | 4 | 2.150 | 1.459 | 0.244 | 0.940 | 0.200 | 1.000 |
| 7 | 3mss | 5 | 1.466 | 1.588 | 0.815 | 0.822 | 0.800 | 0.900 |
| 8 | 3zdg | 4 | 2.964 | 2.873 | 0.651 | 0.700 | 0.600 | 0.600 |
| 9 | 2qnq | 4 | 2.480 | 1.783 | 0.545 | 0.919 | 0.200 | 0.800 |
| 10 | 4lzs | 5 | 0.493 | 0.331 | 0.850 | 0.937 | 0.600 | 0.900 |
| 11 | 1ps3 | 5 | 1.781 | 1.475 | 0.701 | 0.892 | 0.300 | 0.600 |
| 12 | 3l7b | 5 | 1.640 | 1.271 | 0.500 | 0.628 | 0.300 | 0.700 |
| 13 | 2w66 | 5 | 0.868 | 0.938 | 0.857 | 0.624 | 0.800 | 0.700 |
| 14 | 3aru | 5 | 0.889 | 0.853 | 0.836 | 0.956 | 0.700 | 0.900 |
| 15 | 4ih5 | 3 | 1.477 | 1.165 | 0.985 | 0.999 | 1.000 | 1.000 |
| 16 | 1gpk | 5 | 2.285 | 1.472 | -0.517 | 0.921 | -0.900 | 1.000 |
| 18 | 2wbg | 5 | 1.183 | 1.208 | 0.641 | 0.435 | 0.500 | 0.400 |
| 19 | 3udh | 4 | 2.148 | 1.722 | 0.977 | 0.980 | 1.000 | 1.000 |
| 20 | 4jfs | 5 | 1.103 | 1.513 | 0.011 | 0.268 | 0.200 | -0.100 |
| 21 | 3rr4 | 3 | 1.008 | 1.066 | -0.821 | 0.195 | -0.500 | -0.500 |
| 22 | 4dli | 5 | 0.820 | 0.773 | 0.725 | 0.802 | 0.900 | 0.800 |
| 26 | 1q8t | 4 | 1.036 | 0.710 | 0.740 | 0.993 | 0.800 | 1.000 |
| 29 | 4eor | 4 | 0.824 | 0.601 | 0.393 | 0.956 | 0.400 | 0.800 |
| 30 | 4k77 | 5 | 1.115 | 1.567 | 0.927 | 0.894 | 0.600 | 0.800 |
| 31 | 4f09 | 5 | 1.759 | 1.632 | 0.096 | 0.883 | 0.200 | 0.900 |
| 32 | 2brb | 5 | 1.187 | 1.017 | 0.963 | 0.890 | 0.900 | 0.900 |
| 38 | 4jsz | 5 | 1.847 | 1.663 | 0.669 | 0.764 | 0.900 | 0.900 |
| 39 | 2xdl | 4 | 1.991 | 1.539 | 0.751 | 0.922 | 0.800 | 0.800 |
| 40 | 1z95 | 5 | 1.613 | 1.179 | -0.319 | 0.035 | -0.100 | 0.400 |
| 42 | 3g2z | 4 | 1.298 | 1.005 | 0.734 | 0.925 | 0.200 | 1.000 |
| 44 | 4mgd | 5 | 2.187 | 1.884 | 0.683 | 0.347 | 0.700 | 0.600 |
| 45 | 2y5h | 4 | 2.289 | 1.689 | -0.047 | 0.805 | 0.000 | 0.800 |
| 46 | 1nc3 | 5 | 2.915 | 2.835 | 0.548 | 0.732 | 0.200 | 0.700 |
| 48 | 2yge | 5 | 1.289 | 1.204 | -0.124 | 0.249 | -0.200 | 0.500 |
| 49 | 4kzq | 5 | 0.705 | 0.940 | 0.675 | -0.772 | 0.600 | -0.700 |
| 50 | 2ymd | 5 | 2.393 | 2.318 | 0.877 | 0.743 | 0.900 | 0.900 |
| 51 | 3ozt | 5 | 1.576 | 2.293 | 0.473 | 0.781 | 0.500 | 0.800 |
| 54 | 4bkt | 5 | 1.581 | 1.720 | 0.685 | 0.619 | 0.600 | 0.500 |

**Table 4.** Metrics tracked for PCM and AEVPLIG models on various train/test splits of the SAIR-FEP-HID split. 80/85/90/95 indicates the confidence score filter applied to the co-folded data.

| Description | Training set | Testing set | RMSE ↓ [pK Units] | Spearman ↑ | Pearson ↑ |
|---|---|---|---|---|---|
| PCM + Vinardo | SAIR-FEP-HID<sub>train</sub> | SAIR-FEP-HID<sub>test</sub> | 1.260 ± 0.011 | 0.348 ± 0.011 | 0.357 ± 0.011 |
| PCM + Docking + AFEP | SAIR-FEP-HID<sub>train</sub> | SAIR-FEP-HID<sub>test</sub> | 1.257 ± 0.012 | 0.377 ± 0.011 | 0.388 ± 0.011 |
| PCM | SAIR-FEP-HID<sub>train</sub> | SAIR-FEP-HID<sub>test</sub> | 1.198 ± 0.010 | 0.394 ± 0.011 | 0.405 ± 0.011 |
| PCM + Docking + AFEP + Vinardo | SAIR-FEP-HID<sub>train</sub> | SAIR-FEP-HID<sub>test</sub> | 1.225 ± 0.011 | 0.401 ± 0.011 | 0.411 ± 0.011 |
| PCM + AFEP | SAIR-FEP-HID<sub>train</sub> | SAIR-FEP-HID<sub>test</sub> | 1.187 ± 0.010 | 0.410 ± 0.010 | 0.420 ± 0.010 |
| PCM + Docking + Vinardo | SAIR-FEP-HID<sub>train</sub> | SAIR-FEP-HID<sub>test</sub> | 1.211 ± 0.011 | 0.421 ± 0.011 | 0.429 ± 0.010 |
| PCM + Docking | SAIR-FEP-HID<sub>train</sub> | SAIR-FEP-HID<sub>test</sub> | 1.160 ± 0.010 | 0.457 ± 0.010 | 0.461 ± 0.010 |
| AEVPLIG | SAIR-FEP-HID<sub>train</sub> | SAIR-FEP-HID<sub>test</sub> | 1.160 ± 0.010 | 0.457 ± 0.010 | 0.463 ± 0.010 |
| AEVPLIG | SAIR-FEP-HID-60<sub>train</sub> | SAIR-FEP-HID-90<sub>test</sub> | 1.163 ± 0.014 | 0.513 ± 0.014 | 0.518 ± 0.013 |
| AEVPLIG | SAIR-FEP-HID-65<sub>train</sub> | SAIR-FEP-HID-90<sub>test</sub> | 1.159 ± 0.014 | 0.518 ± 0.014 | 0.523 ± 0.013 |
| AEVPLIG | SAIR-FEP-HID-70<sub>train</sub> | SAIR-FEP-HID-90<sub>test</sub> | 1.165 ± 0.014 | 0.510 ± 0.014 | 0.515 ± 0.013 |
| AEVPLIG | SAIR-FEP-HID-75<sub>train</sub> | SAIR-FEP-HID-90<sub>test</sub> | 1.155 ± 0.014 | 0.517 ± 0.014 | 0.524 ± 0.013 |
| AEVPLIG | SAIR-FEP-HID-80<sub>train</sub> | SAIR-FEP-HID-90<sub>test</sub> | 1.172 ± 0.014 | 0.500 ± 0.014 | 0.507 ± 0.013 |
| AEVPLIG | SAIR-FEP-HID-85<sub>train</sub> | SAIR-FEP-HID-90<sub>test</sub> | 1.156 ± 0.014 | 0.514 ± 0.014 | 0.518 ± 0.013 |
| AEVPLIG | SAIR-FEP-HID-90<sub>train</sub> | SAIR-FEP-HID-90<sub>test</sub> | 1.177 ± 0.014 | 0.484 ± 0.014 | 0.487 ± 0.014 |
| AEVPLIG | SAIR-FEP-HID-95<sub>train</sub> | SAIR-FEP-HID-90<sub>test</sub> | 1.263 ± 0.014 | 0.305 ± 0.016 | 0.333 ± 0.015 |
| AEVPLIG | SAIR-FEP-HID-60<sub>train</sub> | SAIR-FEP-HID-95<sub>test</sub> | 1.103 ± 0.032 | 0.547 ± 0.033 | 0.568 ± 0.030 |
| AEVPLIG | SAIR-FEP-HID-65<sub>train</sub> | SAIR-FEP-HID-95<sub>test</sub> | 1.096 ± 0.031 | 0.557 ± 0.033 | 0.571 ± 0.030 |
| AEVPLIG | SAIR-FEP-HID-70<sub>train</sub> | SAIR-FEP-HID-95<sub>test</sub> | 1.079 ± 0.030 | 0.572 ± 0.032 | 0.588 ± 0.029 |
| AEVPLIG | SAIR-FEP-HID-75<sub>train</sub> | SAIR-FEP-HID-95<sub>test</sub> | 1.072 ± 0.029 | 0.574 ± 0.031 | 0.591 ± 0.029 |
| AEVPLIG | SAIR-FEP-HID-80<sub>train</sub> | SAIR-FEP-HID-95<sub>test</sub> | 1.072 ± 0.029 | 0.573 ± 0.032 | 0.589 ± 0.030 |
| AEVPLIG | SAIR-FEP-HID-85<sub>train</sub> | SAIR-FEP-HID-95<sub>test</sub> | 1.067 ± 0.029 | 0.588 ± 0.031 | 0.602 ± 0.029 |
| AEVPLIG | SAIR-FEP-HID-90<sub>train</sub> | SAIR-FEP-HID-95<sub>test</sub> | 1.076 ± 0.031 | 0.569 ± 0.034 | 0.588 ± 0.032 |
| AEVPLIG | SAIR-FEP-HID-95<sub>train</sub> | SAIR-FEP-HID-95<sub>test</sub> | 1.145 ± 0.035 | 0.437 ± 0.036 | 0.468 ± 0.035 |

**Table 5.** Metrics for AEVPLIG models trained on various splits of the SAIR-OOD data split. 80/85/90/95 indicates the confidence score filter applied to the co-folded data.

| Test Dataset | Train Dataset | Model | RMSE ↓ [pK Units] | Pearson ↑ | Kendall ↑ | Spearman ↑ |
|---|---|---|---|---|---|---|
| CASF v2016 | Refined2020++ | AEVPLIG | 1.745 ± 0.160 | 0.669 ± 0.072 | 0.472 ± 0.077 | 0.657 ± 0.091 |
| CASF v2016 | Refined2020++ | PCM | 1.690 ± 0.160 | 0.690 ± 0.072 | - | 0.680 ± 0.080 |
| AEVPLIG OOD | Refined2020+ | AEVPLIG | 1.514 ± 0.142 | 0.692 ± 0.063 | 0.514 ± 0.050 | 0.711 ± 0.058 |
| AEVPLIG OOD | Refined2020+ | PCM | 1.800 ± 0.110 | 0.490 ± 0.068 | - | 0.520 ± 0.080 |
| CASF v2016 | SAIR-OOD-95 + Refined2020+ | AEVPLIG | 1.713 ± 0.149 | 0.684 ± 0.070 | 0.482 ± 0.068 | 0.669 ± 0.080 |
| AEVPLIG OOD | SAIR-OOD-95 + Refined2020+ | AEVPLIG | 1.505 ± 0.129 | 0.690 ± 0.058 | 0.501 ± 0.053 | 0.697 ± 0.059 |
| CASF v2016 | SAIR-OOD-90 + Refined2020++ | AEVPLIG | 1.642 ± 0.154 | 0.708 ± 0.076 | 0.512 ± 0.074 | 0.694 ± 0.037 |
| CASF v2016 | SAIR-OOD-90 + Refined2020++ | PCM | 1.690 ± 0.140 | 0.690± 0.060 | - | 0.700 ± 0.060 |
| AEVPLIG OOD | SAIR-OOD-90 + Refined2020+ | AEVPLIG | 1.471 ± 0.123 | 0.709 ± 0.054 | 0.513 ± 0.052 | 0.711 ± 0.056 |
| AEVPLIG OOD | SAIR-OOD-90 + Refined2020+ | PCM | 1.800 ± 0.110 | 0.510 ± 0.080 | - | 0.520 ± 0.100 |
| CASF v2016 | SAIR-OOD-85 + Refined2020++ | AEVPLIG | **1.607 ± 0.146** | **0.727 ± 0.072** | **0.523 ± 0.077** | **0.706 ± 0.087** |
| AEVPLIG OOD | SAIR-OOD-85 + Refined2020+ | AEVPLIG | 1.515 ± 0.131 | 0.683 ± 0.055 | 0.506 ± 0.051 | 0.703 ± 0.058 |
| CASF v2016 | SAIR-OOD-80 + Refined2020++ | AEVPLIG | 1.722 ± 0.173 | 0.647 ± 0.079 | 0.446 ± 0.074 | 0.618 ± 0.092 |
| AEVPLIG OOD | SAIR-OOD-80 + Refined2020+ | AEVPLIG | 1.495 ± 0.149 | 0.698 ± 0.061 | 0.522 ± 0.053 | 0.718 ± 0.061 |
| CASF v2016 | SAIR-OOD + Refined2020++ | PCM | 1.840 ± 0.180 | 0.630 ± 0.070 | - | 0.640 ± 0.080 |
| AEVPLIG OOD | SAIR-OOD + Refined2020+ | AEVPLIG | 1.476 ± 0.126 | 0.712 ± 0.059 | 0.527 ± 0.056 | 0.719 ± 0.063 |
| AEVPLIG OOD | SAIR-OOD + Refined2020+ | PCM | 1.780 ± 0.120 | 0.580 ± 0.080 | - | 0.520 ± 0.100 |
| CASF v2016 | 20% SAIR-OOD-90 + Refined2020++ | AEVPLIG | 1.686 ± 0.158 | 0.694 ± 0.075 | 0.501 ± 0.074 | 0.677 ± 0.088 |
| AEVPLIG OOD | 20% SAIR-OOD-90 + Refined2020+ | AEVPLIG | 1.544 ± 0.136 | 0.676 ± 0.056 | 0.501 ± 0.051 | 0.695 ± 0.058 |
| CASF v2016 | 40% SAIR-OOD-90 + Refined2020++ | AEVPLIG | 1.672 ± 0.158 | 0.697 ± 0.074 | 0.496 ± 0.074 | 0.678 ± 0.087 |
| AEVPLIG OOD | 40% SAIR-OOD-90 + Refined2020+ | AEVPLIG | 1.472 ± 0.128 | 0.702 ± 0.059 | 0.524± 0.054 | 0.721± 0.061 |
| CASF v2016 | 60% SAIR-OOD-90 + Refined2020++ | AEVPLIG | 1.645 ± 0.176 | 0.703 ± 0.079 | 0.506 ± 0.076 | 0.685 ± 0.091 |
| AEVPLIG OOD | 60% SAIR-OOD-90 + Refined2020+ | AEVPLIG | 1.481 ± 0.124 | 0.705± 0.057 | 0.521 ± 0.055 | 0.713 ± 0.062 |
| CASF v2016 | 80% SAIR-OOD-90 + Refined2020++ | AEVPLIG | 1.666 ± 0.157 | 0.698 ± 0.072 | 0.505 ± 0.073 | 0.687 ± 0.085 |
| AEVPLIG OOD | 80% SAIR-OOD-90 + Refined2020+ | AEVPLIG | **1.459± 0.121** | **0.715± 0.053** | **0.540 ± 0.049** | **0.738 ± 0.054** |

**Table 6.**
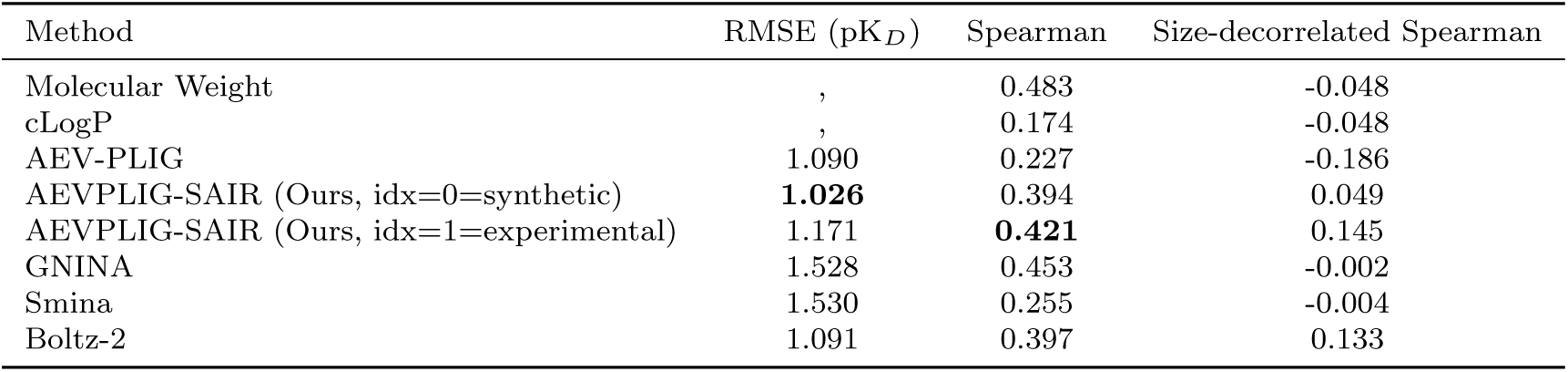
Affinity prediction on the OpenBind EV-A71 2A protease benchmark (*n* = 494 compounds). The size-decorrelated Spearman is the partial Spearman correlation between predictions and experimental affinity while controlling for heavy-atom count, *ρ*(pred, aff | HAC); it removes the molecular-size confounder intrinsic to this fragment-elaboration series.

| Method | RMSE (pK<sub>*D*</sub>) | Spearman | Size-decorrelated Spearman |
|---|---|---|---|
| Molecular Weight | , | 0.483 | -0.048 |
| cLogP | , | 0.174 | -0.048 |
| AEV-PLIG | 1.090 | 0.227 | -0.186 |
| AEVPLIG-SAIR (Ours, idx=0=synthetic) | **1.026** | 0.394 | 0.049 |
| AEVPLIG-SAIR (Ours, idx=1=experimental) | 1.171 | **0.421** | 0.145 |
| GNINA | 1.528 | 0.453 | -0.002 |
| Smina | 1.530 | 0.255 | -0.004 |
| Boltz-2 | 1.091 | 0.397 | 0.133 |

**Table 7.** Benchmarking AEV-PLIG on GEMS’s leakage-free PDBbind CleanSplit [14], evaluated on CASF-2016 (285 complexes; “indep.” is the 144-complex leakage-independent subset). Trained on CleanSplit alone, AEV-PLIG generalizes competitively; adding SAIR synthetic data (dual-source, leakage-filtered against CASF) improves performance further on both the full set and the leakage-independent subset (0.03 to 0.04 Pearson), showing that the benefit of synthetic augmentation is not a train/test-leakage artifact. The Pearson/RMSE reported for GEMS, GenScore, and Pafnucy are on the full CASF-2016 set as given in [14]; GEMS additionally reports r.m.s.e. 1.367 on the independent subset (Pearson not reported), hence the “, ” entries.

| Model (trained on PDBbind CleanSplit) | Pearson | RMSE (pK) | Pearson (indep.) |
|---|---|---|---|
| AEV-PLIG (experimental only) | 0.760 | 1.446 | 0.759 |
| AEV-PLIG + SAIR (dual-source) | **0.793** | **1.403** | **0.802** |
| GEMS | 0.803 | 1.308 | , |
| GenScore | 0.780 | 1.362 | , |
| Pafnucy | 0.746 | 1.484 | , |

**Table 8.**
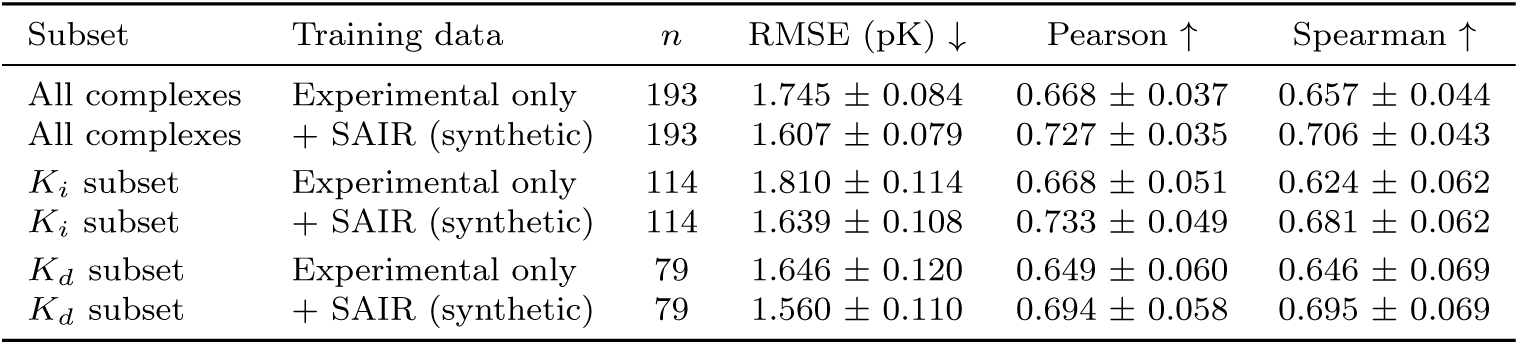
Effect of SAIR (synthetic) augmentation broken down by experimental measurement type on the CASF-2016 test set. The SAIR augmentation data is composed entirely of IC_50_ measurements, whereas CASF-2016 contains only *K_i_* (114 complexes) and *K_d_*(79 complexes) values and no IC_50_. We compare the same AEVPLIG architecture trained on experimental data alone versus experimental plus SAIR synthetic data, evaluated on each measurement-type subset. Augmentation improves accuracy on both the *K_i_*and *K_d_* subsets even though the synthetic data contains neither measurement type, indicating that the added signal is not specific to the IC_50_ label. Uncertainties are *±*1 s.d. from a 1000-iteration bootstrap over complexes.

| Subset | Training data | *n* | RMSE (pK) ↓ | Pearson ↑ | Spearman ↑ |
|---|---|---|---|---|---|
| All complexes | Experimental only | 193 | 1.745 ± 0.084 | 0.668 ± 0.037 | 0.657 ± 0.044 |
| All complexes | + SAIR (synthetic) | 193 | 1.607 ± 0.079 | 0.727 ± 0.035 | 0.706 ± 0.043 |
| *K<sub>i</sub>* subset | Experimental only | 114 | 1.810 ± 0.114 | 0.668 ± 0.051 | 0.624 ± 0.062 |
| *K<sub>i</sub>* subset | + SAIR (synthetic) | 114 | 1.639 ± 0.108 | 0.733 ± 0.049 | 0.681 ± 0.062 |
| *K<sub>d</sub>* subset | Experimental only | 79 | 1.646 ± 0.120 | 0.649 ± 0.060 | 0.646 ± 0.069 |
| *K<sub>d</sub>* subset | + SAIR (synthetic) | 79 | 1.560 ± 0.110 | 0.694 ± 0.058 | 0.695 ± 0.069 |

**Table 9.**
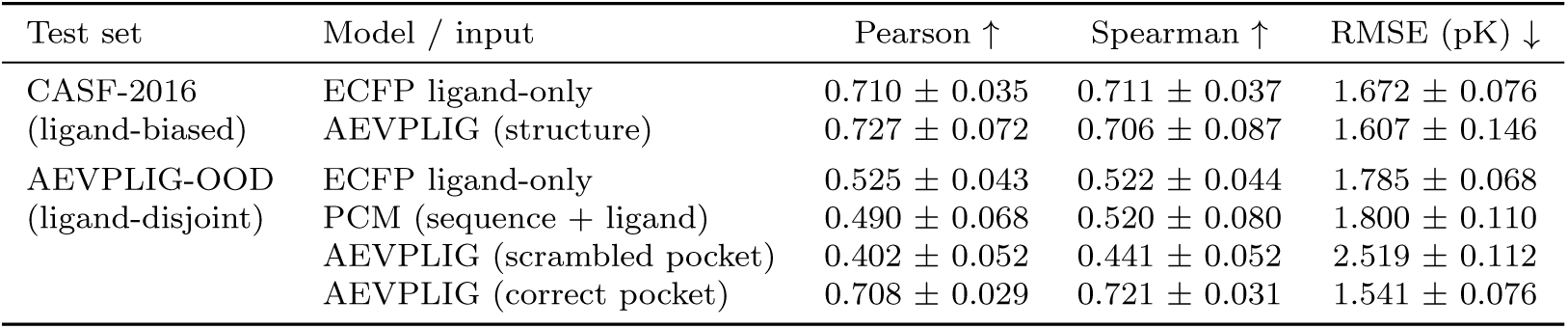
Ligand-only baseline and pocket-scrambled control probing whether AEVPLIG uses the 3D protein–ligand structure or merely ligand identity (addressing the ligand-memorization concern of Ref. [19]). On the CASF-2016 core set, whose ligands have close analogues in the training data and which is therefore strongly ligand-biased, an ECFP-fingerprint ligand-only model (gradient boosting on 2048-bit Morgan fingerprints) matches the structure-based AEVPLIG, showing that standard benchmarks cannot distinguish genuine structure use from ligand memorization. On the leakage-filtered, ligand-disjoint AEVPLIG-OOD set (only 1.4% of test ligands appear in training), both non-structural baselines (ECFP ligand-only and the sequence+ligand PCM model) collapse, and scrambling the pocket, pairing each ligand with a random incorrect protein, degrades AEVPLIG to a similar level, whereas AEVPLIG with the correct pocket retains strong performance. This demonstrates that on tasks requiring generalization the model genuinely relies on the 3D structure. Uncertainties are *±*1 s.d. from a 1000-iteration bootstrap; the ligand-only/scramble rows use *n* = 266 OOD complexes with locally available structures.

| Test set | Model / input | Pearson ↑ | Spearman ↑ | RMSE (pK) ↓ |
|---|---|---|---|---|
| CASF-2016 | ECFP ligand-only | 0.710 ± 0.035 | 0.711 ± 0.037 | 1.672 ± 0.076 |
| (ligand-biased) | AEVPLIG (structure) | 0.727 ± 0.072 | 0.706 ± 0.087 | 1.607 ± 0.146 |
| AEVPLIG-OOD | ECFP ligand-only | 0.525 ± 0.043 | 0.522 ± 0.044 | 1.785 ± 0.068 |
| (ligand-disjoint) | PCM (sequence + ligand) | 0.490 ± 0.068 | 0.520 ± 0.080 | 1.800 ± 0.110 |
|  | AEVPLIG (scrambled pocket) | 0.402 ± 0.052 | 0.441 ± 0.052 | 2.519 ± 0.112 |
|  | AEVPLIG (correct pocket) | 0.708 ± 0.029 | 0.721 ± 0.031 | 1.541 ± 0.076 |

To characterise when synthetic augmentation helps, we stratified the change in AEVPLIG performance (the same architecture trained with versus without SAIR data, evaluated on CASF 2016 and AEVPLIG-OOD) by affinity, target, and ligand chemotype (Figure 6). The benefit is concentrated at the extremes of the affinity range (Figure 6a): the experimental-only model regresses strongly toward the training mean (predicted pK spanning a standard deviation of 1.0 against a true value of 2.2), and augmentation widens this range, reducing RMSE most for the weakest (pK *<* 4) and most potent (pK *>* 9) binders while being neutral in the data-dense mid-range (pK 4–6). Across CASF 2016 targets, the gains are largest precisely where the experimental-only model ranks poorly: the per-target improvement in Pearson correlation is inversely related to the baseline model’s per-target performance (Spear-man *ρ* = *−*0.73; Figure 6b), while targets already well described by crystal data are essentially unchanged. Finally, augmentation is chemotype-agnostic on the in-domain crystal test but becomes selective out of distribution (Figure 6c), where it preferentially benefits smaller, more aliphatic ligands and can modestly degrade predictions for large, poly-aromatic, lipophilic molecules, consistent with the co-folded poses of such ligands being less reliable. Together, these trends indicate that synthetic augmentation is most valuable at the affinity extremes and for the targets and chemotypes under-represented in the experimental data, and is largely neutral where experimental coverage is already dense.

**Fig. 5.**
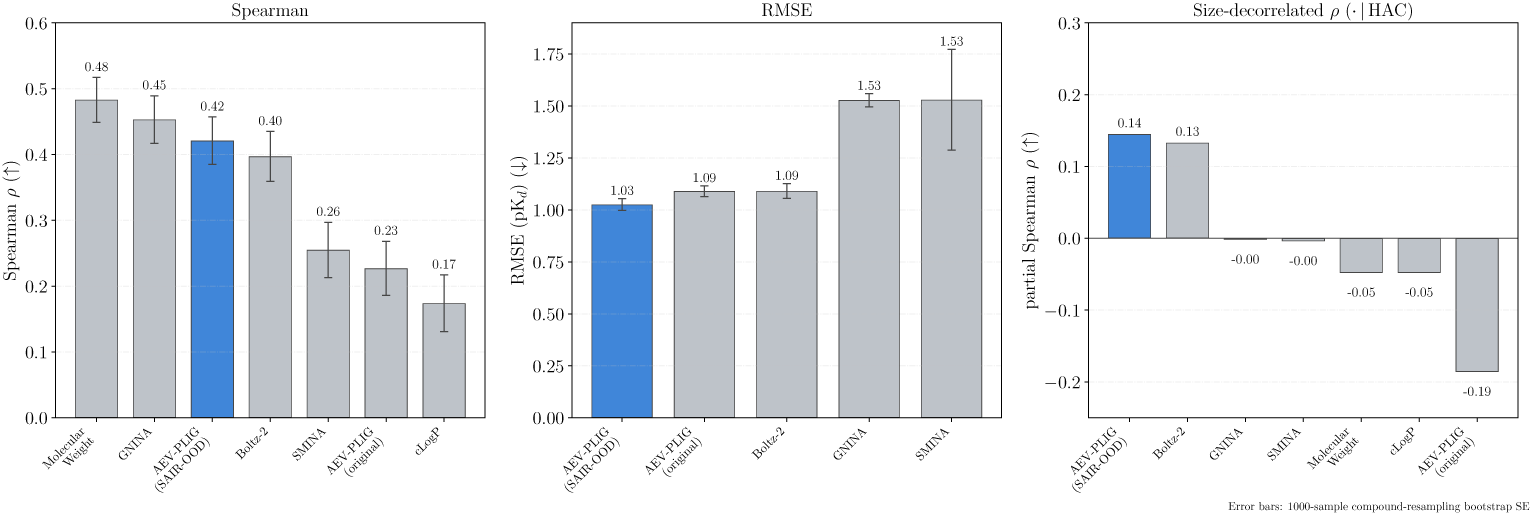
Performance of various methods on the OpenBind EV-A71 2A protease benchmark. Left: Spearman rank correlation between predicted and experimental affinity across compounds, without any correction for molecular size. Centre: root mean squared error (RMSE). Right: size-decorrelated ranking, the partial Spearman correlation between prediction and experimental affinity while controlling for heavy-atom count (HAC), *ρ*(pred, aff | HAC), which removes the molecular-size confound present in this fragment-elaboration series. The AEVPLIG model trained on SAIR and Refined2020+(blue) achieves the lowest RMSE and the highest size-decorrelated ranking; under the uncorrected Spearman metric it is on par with the docking and molecular-size baselines (grey), whose ranking is inflated by their correlation with molecular size. Error bars are the standard error from a 1000-iteration compound-resampling bootstrap.

**Fig. 6.**
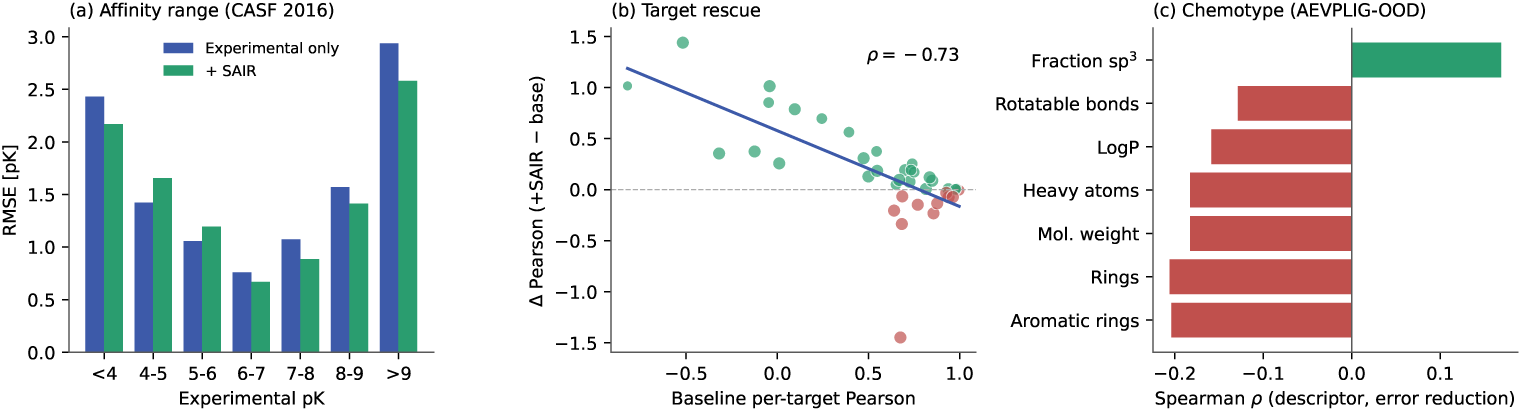
When does synthetic (SAIR) augmentation help? The same AEVPLIG architecture is trained with and without SAIR data and evaluated on identical held-out complexes. (a) RMSE by experimental affinity bin on CASF 2016: augmentation reduces error most at the affinity extremes and is neutral in the data-dense mid-range. (b) Per-target improvement in Pearson correlation versus the baseline model’s per-target performance across CASF 2016 targets (point area *∝* number of complexes); gains are largest where the experimental-only model ranks poorly (Spearman *ρ* = *−*0.73). (c) Spearman correlation between ligand descriptors and the per-complex reduction in absolute error on AEVPLIG-OOD: augmentation preferentially benefits smaller, more aliphatic ligands and degrades large, poly-aromatic, lipophilic ones.

#### 3.2.2 Neural Scaling with high-quality structural data in the low data regime

To understand a model’s scaling performance, one typically assesses its test-set performance as the number of training points increases. For structure-based deep learning models trained on synthetic protein-ligand complexes, this and previous work suggest that naively increasing the dataset size does not lead to performance improvements [21]. However, given that our other analyses revealed that the quality of the structure data is a key variable in model performance, we sought to replicate this pattern in our data and evaluate whether adjusting the quality of structures could recover this finding. To evaluate this, we train models with different dataset sizes by changing the confidence score filter (see Table 1). We then monitor the performance of AEVPLIG on CASF 2016 and AEVPLIG-OOD. Compared with the experimental baseline, models trained with more data also show performance gains on the test sets. However, similar to what is observed in Figure 2b, the metrics do not continue to improve as the number of complexes increases.

Since adding low-confidence structures to the data does not yield typical model scaling behaviour, we evaluated whether improving the quality of structures in the training set could help recover it. To do so, we randomly sample 20%, 40%, 60%, and 80% of SAIR-OOD-90, and train on the selected samples along with the Refined2020+ and Refined2020++ data. Next, we evaluate the performance of the AEVPLIG model on CASF 2016 and AEVPLIG-OOD as a function of dataset size. Applying this quality filter allows us to roughly recover typical model scaling behaviour as seen in Figure 4b. When comparing the differences in the curves between CASF 2016 and AEVPLIG-OOD, the curves for AEVPLIG-OOD have slopes with magnitudes 3-6 times larger than those of CASF 2016. This indicates that the data aggregation strategy used to construct SAIR [25] may exhibit stronger overlap with the complexes in AEVPLIG-OOD than with those in CASF 2016. From these scaling curves, one can estimate how many complexes are needed to observe a particular metric value. As an example, to achieve a Pearson correlation coefficient of 0.9 for AEVPLIG-OOD would require ≈ 32 ×10^6^ complexes. For CASF 2016, this many complexes would yield a Pearson correlation of 0.723, indicating, once again, that the data aggregation strategy used in SAIR is currently inadequate for this test split. Taken together, these results suggest that by including only high-quality co-folded structures during training, we can recover traditional model scaling behaviour. However, further experimentation with larger datasets is needed to confirm these findings. This paves the way for large-scale computational data augmentation to build affinity prediction models that perform effectively on extreme generalization tasks.

#### 3.2.3 Performance on the EV-A71 2A Protease target from OpenBind

To further understand the generalizability and applicability of the models trained in this work, we ran inference on the recent OpenBind affinity prediction benchmark [42] using the AEVPLIG model trained on SAIR-OOD and Refined2020+. The OpenBind EV-A71 2A protease benchmark is a dense, single-target structure–affinity dataset released by the OpenBind Consortium. The dataset contains 925 crystallographic protein–ligand binding events across 699 compounds, of which 494 have experimental affinity measurements derived from filtered dose–response data. Ground-truth structures were determined by X-ray crystallography at the Diamond Light Source. To confirm that this benchmark is genuinely out-of-distribution with respect to our training data, we quantified its similarity to the training set: the closest training protein shares only 47% sequence identity with the EV-A71 2A protease (a single full-domain homolog), and the most similar training ligand has a Tanimoto similarity of 0.74, with no identical ligands present. This confirms that neither the target nor its chemotypes are represented in SAIR/SAIR-OOD training.

Following the official evaluation protocol, predictions were made for all 649 structures with paired crystallographic coordinates and affinity labels. Compound-level metrics were computed by grouping structures by canonical SMILES and averaging predictions for compounds represented by multiple crystal structures. Performance was assessed using RMSE and Spearman rank correlation coefficient, computed over 494 unique compounds. We evaluated the model twice, once with the source embedding for synthetic data and once with the source embedding for experimental data. As shown in Figure 5, we find that our models yield the lowest RMSE when compared to the physics-based and AI-based methods included in the current benchmark. Interestingly, this result was found when using the synthetic data embedding. In addition, we find that, among the AI-based methods, that our AEVPLIG model (with the experimental source embedding) had the highest Spearman ranking coefficient. We see a 1.85x improvement when compared to the original AEVPLIG model training on experimental data alone. Under the uncorrected Spearman metric the new AEVPLIG model is on-par with gnina and does not outperform molecular weight; however, this ranking is confounded by molecular size, as we discuss next.

Experimental affinity in the EV-A71 2A fragment-elaboration series is strongly coupled to molecular size, correlating with heavy-atom count (HAC) at *ρ* = 0.53. To isolate genuine affinity ranking from this confounder, we additionally report the partial Spearman correlation between predictions and experimental affinity while controlling for HAC, *ρ*(pred, aff HAC) (Table 6). This is computed by rank-transforming the predictions, experimental affinity, and heavy-atom count, regressing the heavy-atom-count rank out of both the prediction and affinity ranks, and correlating the two sets of residuals. Under this size-decorrelated metric, the apparent performance of size-driven methods collapses: molecular weight, cLogP, GNINA, Smina, and the baseline AEV-PLIG all fall to near-zero or negative values, with the molecular-weight and GNINA predictions themselves correlating with HAC at 0.94 and 0.86, respectively. Only our SAIR-augmented model (0.145) and Boltz-2 (0.133) retain a meaningful size-independent ranking signal, with our model achieving the highest value. We therefore refrain from claiming superiority under the uncorrected Spearman metric, but emphasize that once size effects are removed, our approach is the strongest ranker on this benchmark. Applying the same decorrelation to the AEVPLIG-OOD test set leaves our Spearman essentially unchanged (0.575), indicating that size is not a confounder there.

## 4 Conclusion

In this work, we investigate the role of incorporating synthetic data in improving deep-learning-based experimental binding affinity predictions for protein-ligand complexes. We first extend the SAIR dataset by adding additional physics-based data. We also create two additional data splits to aid future researchers in benchmarking deep-learning affinity models. Next, we demonstrate that augmenting experimentally derived structural data with synthetic data, generated by a co-folding model, can improve downstream model performance on challenging out-of-distribution prediction tasks.

We initially create a core dataset, SAIR-FEP, in which we perform docking and AFEP calculations for ≈ 80, 000 complexes and split the data into train/test sets to simulate realistic drug discovery scenarios. Next, we curate a train/test split, called SAIR-FEP-HID, which is intended to be a true out-of-domain split in which the target is not included in the training set. With this set, we train PCM models and show that including additional physics-based features (i.e., docking scores and AFEP values) improves test-set performance. We also compared the PCM models to a DL approach, AEVPLIG, which demonstrates competitive performance alongside the highest-performing physics-augmented PCM variant. However, introducing more co-folded complexes with lower confidence scores into AEVPLIG does not improve test-set metrics, contrary to the typical scaling laws observed in deep learning models.

The second train/test split of SAIR-FEP, called SAIR-FEP-OPT, is intended to simulate a lead-optimization scenario. Here, the training and testing sets have the same targets but different ligands. With this split, we compare the correlation of models trained independently for each protein to that of a model trained across all proteins. Among the global models, the PCM base model achieved the highest mean Pearson correlation. It notably outperformed variants that incorporated additional physics-based features such as docking and AFEP values.

We also curate SAIR-OOD, an additional split in which we remove SAIR entries with similar complexes from the CASF 2016 and AEVPLIG-OOD test sets to prevent data leakage. We then train AEVPLIG models on synthetic and experimental data simultaneously and show that including synthetic data improves test set metrics on the experimental data. For the OpenBind benchmark, our models yield the lowest root mean squared errors and a 1.85x improvement of Spearman ranking correlation when compared to the original AEVPLIG model training on experimental data alone. We also investigate model performance as more, lower-confidence co-folded structures are added to the training set and again observe metric saturation. To test if typical scaling laws are observed, we randomly select entries from the high-confidence complexes in increasing amounts and monitor the test metrics. Here, we find that adding more complexes improves performance, indicating typical scaling-law behaviour of DL models for binding affinity prediction.

This work demonstrates that including synthetic data in various ways generally improves the predictive performance of models for experimental binding affinities. However, we show that one cannot blindly include synthetic data in their training. To achieve performance improvements, careful curation of the co-folded structures is required. Taken together, these results show that by including only high-quality cofolded structures during training, we can recover traditional model scaling behaviour. This paves the way for large-scale computational data augmentation efforts to build affinity prediction models that perform effectively on extreme generalization tasks common in drug discovery.

## 5 Data Availability

The docking and absolute free energy values for both SAIR and PDBBind can be found in a comma separated value format in the Github repository found here: https://github.com/sandbox-quantum/on-improving-experimental-binding-affinity-predictions-with-synthetic-data. The AEVPLIG-OOD train/test split is found here: https://github.com/isakvals/OOD-Test. The structures from SAIR can be down-loaded from Hugging Face: https://huggingface.co/datasets/SandboxAQ/SAIR and the structures from PDBBind (including CASF 2016) can be downloaded from here: https://www.pdbbind-plus.org.cn/.

## 6 Code Availability

To train the AEVPLIG models, we used the following Github repository: https://github.com/weitse-hsu/AEV-PLIG-refined. A patch can be applied for our model by using the code found here: https://github.com/sandbox-quantum/on-improving-experimental-binding-affinity-predictions-with-synthetic-data. Scripts to train proteochemetric and XGBoost models can also be found in this repo.

## 7 Author Contributions

K. R., P. P. Z. and A. B. prepared data, trained models, wrote and reviewed the manuscript. J. C-D. advised, wrote, and reviewed the manuscript, L. L and P. K. J. contributed the docking and AFEP calculations, wrote and reviewed the manuscript. H. R., L. B., J. O., B. J. S., P. L., S. B. and M. G. advised and reviewed the manuscript. M. v. D. contributed to data preparation, advised, and reviewed the manuscript. A. S., L. H., and M. P. developed the AFEP method, advised, and reviewed the manuscript.

## A Stratified analysis of when synthetic augmentation helps

## B Per-target significance testing

To test whether the improvements from synthetic augmentation are statistically significant rather than aggregate point estimates, we computed each metric per target and compared the experimental-only baseline against the SAIR-augmented model across targets using a paired Wilcoxon signed-rank test (Table 2). For CASF 2016 the per-target grouping is the target clustering supplied with the PDBbind/CASF 2016 core set, in which the 285 core complexes are assigned to 57 protein targets by 90% sequence similarity (five complexes per target); this grouping differs from the *N*_proteins_ = 156 entry of Table 1, which counts distinct protein sequences, because a single target contains several point mutants or constructs with near-identical but non-identical sequences. For AEVPLIG-OOD, which has no such supplied clustering, targets are defined by distinct protein sequence. Only targets with at least three complexes are included in the test: this retains 38 of the 49 CASF 2016 clusters present in our evaluation (176 of the 193 complexes; the remaining 17 fall in 11 clusters of one or two complexes that are too small to score within a target) and 22 targets on AEVPLIG-OOD. The pooled metrics reported elsewhere use all complexes. Per-target assignments and paired values for every included CASF 2016 cluster are listed in Table 3. On CASF 2016 all four metrics improve significantly (*p <* 0.01); on AEVPLIG-OOD, where per-target estimates are sparse, the models are statistically indistinguishable.

## C Results in Tabular Format

### D How modest improvements in correlation lead to significant improvement for enrichment factors

To begin, we assume there are a certain number of actives, *N*_actives_, from a library of molecules with size *N*_total_. The true and predicted affinities, *y* and *ŷ*, are connected via a standard bi-variate normal distribution function, Φ_2_(*y, ŷ*; *ρ*), where *ρ* is the correlation between *y* and *ŷ*. To find the number of hits, we can integrate this function, i.e.

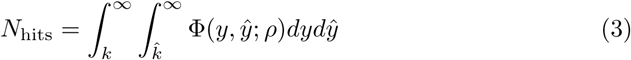

where *k, ^k*, change depending on the threshold taken. Typical choices include 0.01%, 0.1%, 1%, 5% which correspond to values of *k, ^k* = 3.719, 3.090, 2.326, 1.645. These are the number of standard deviations above the mean that a true/predicted structure must exceed to be included in the cutoff. In Table 10, we compute the number of hits found from a library of *N*_total_ = 1, 000, 000 and *N*_actives_ = 1000, taking *ρ* = 0.669 and *ρ* = 0.727, which are the aggregate Pearson correlations of the experimental-only and SAIR-augmented AEVPLIG models on CASF 2016 (the Spearman correlation improves comparably, 0.657 0.706). In a realistic setting, we would likely downselect to 1000 molecules (0.01% cutoff), and in this scenario we see a 41% improvement in the estimated number of hits found (27*→*38).

**Table 10.**
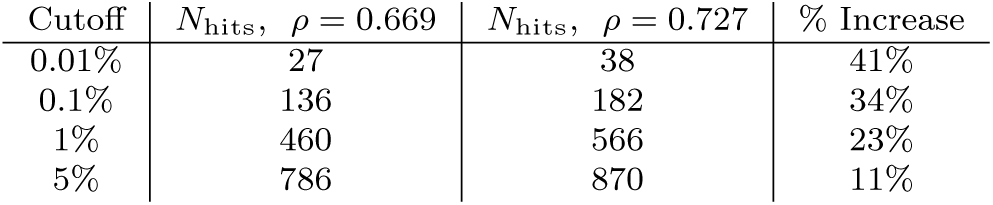
Enrichment performance gains when correlation improves by *≈* 10%, which is what is observed in this manuscript. The total number of compounds *N*_total_ = 1, 000, 000 and the total number of true actives, *N*actives = 1000.

